# Modelling the deathbed of ASF-infected wild boars in South Korea using 2019-2020 national surveillance data

**DOI:** 10.1101/2021.01.26.428275

**Authors:** Jun-Sik Lim, Timothée Vergne, Son-Il Pak, Eutteum Kim

## Abstract

In September 2019, African swine fever (ASF) was reported in South Korea for the first time. Since then, more than 651 ASF cases in wild boars and 14 farm outbreaks have been notified in the country. The purpose of this study was to characterize the spatial distribution of ASF-positive wild boar carcasses to identify the risk factors associated with the presence of ASF and number of ASF-positive wild boar carcasses in the affected areas. To achieve this objective, we divided the study into two periods (October 2, 2019, to January 19, 2020, and January 19 to April 28, 2020) and aggregated the number of reported ASF-positive carcasses into a regular grid of hexagons. To account for imperfect detection, we adjusted spatial zero-inflated Poisson regression models to the number of ASF-positive wild boar carcasses per hexagons. During the first study period, only proximity to North Korea was identified as a risk factor for the presence of African swine fever virus (ASFV). In addition, there were more reports in the affected hexagons with a high habitat suitability for wild boar, low heat load index (HLI), and high human density. During the second study period, proximity to an ASF-positive carcass reported during the first period was the only significant risk factor for the presence of ASF-positive carcasses. Additionally, high HLI and low elevation were associated with an increased number of ASF-positive carcasses reported in the affected hexagons. Although the proportion of ASF-affected hexagons increased from 0.06 (95% credible interval [CrI]: 0.05-0.07) to 0.09 (95% CrI: 0.08-0.10), the probability of reporting ASF-affected hexagons increased substantially from 0.49 (95% CrI: 0.41-0.57) to 0.73 (95% CrI: 0.66-0.81) between the two study periods. These results can be used to further advance risk-based surveillance.

## INTRODUCTION

African swine fever (ASF), caused by the African swine fever virus (ASFV), is a highly contagious viral disease that affects both domestic and wild pigs. Symptoms of ASFV infection include fever, hemorrhage, vomiting and diarrhea, and nearly 100 % mortality can occur with some strains, including that circulating in the Republic of Korea (Kim et al., 2020). Owing to the absence of treatment and vaccine, ASF has imposed a significant socioeconomic burden on the livestock sector and caused a negative impact on the environment (Chenais et al., 2019).

Since ASFV can spill over between wild boars and domestic pigs, it is crucial to understand the disease dynamics in both species. Therefore, government agencies have developed surveillance systems for wild boars. Since ASF has a short infectious period due to severe clinical symptoms, which rapidly leads to death (Gabriel et al., 2011), contact with ASF-positive carcasses or contaminated environment, and consumption of the ASF-positive carcass by scavengers are considered to be the main drivers for the prevalence of ASF among wild boar populations. Thus, one of the most effective strategies for ASF control among wild boars is the detection and disposal of ASF-positive carcasses and disinfection of the surrounding contaminated environment (Probst et al., 2017; O’Neill et al., 2020; Pepin et al., 2020). However, in East and Southeast Asia, wild boar surveillance has not been conducted homogeneously across regions, with many countries having reported only outbreaks in farms.(Vergne et al., 2020). Because diseased animals behave differently from healthy animals, it is possible to infer the deathbed of ASF-infected wild boars, which can be helpful for improving risk-based surveillance strategies and optimizing interventions (Morelle et al., 2019).

In the Republic of Korea, the first case of ASF was reported in a domestic pig farm on September 16, 2019. To mitigate the spread of ASF, the government culled pigs from 261 pig farms in the affected counties (Ministry of Agriculture Food and Rural Affairs, 2020). By early October, only 14 cases of ASF in domestic farms were reported and the epidemic was declared to be over (Yoon et al., 2020). However, according to the passive surveillance system that was already launched before this ASF outbreak, the first case of ASF was reported in a wild boar carcass on October 02, 2019. Later, in November 2019, the government expanded the surveillance system into an enhanced (active and passive) system to include activities such as hunting, trapping, and reporting of carcasses. Despite this effort, the number of reported cases increased in January 2020; these were mainly from wild boar carcasses. Most wildlife surveillance systems have limited detection sensitivity due to dependency on public reporting and limited resources. Therefore, it cannot be determined whether this increase in reports was primarily because of increased surveillance sensitivity or increased prevalence of ASF. Thus, it was difficult to evaluate the dynamics of ASF spread among the wild boar population. Because of the uncertainty of the risk of ASF transmission, as of August 2020, restocking of pigs into the farms that were stamped out in the affected regions is still prohibited. Furthermore, it is challenging to identify the risk factors when the detection probability of a disease is heterogeneous, as this can lead to reporting bias.

Zero-inflated models have been applied in veterinary epidemiological research to identify risk factors of animal disease, while adjusting for imperfect detection (Vergne et al., 2014; Vergne et al., 2016). Zero-inflated count models divide the zeros into true and false zeros at the epidemiological unit level; the former represents true negatives, i.e. disease-free units with zero case reports, and the latter indicates false negatives, i.e. affected units with zero case reports. Thus, the model can be utilized for the estimation of the number of false negative units and therefore of the sensitivity of the surveillance and to clarify the actual risk of a disease by accounting for reporting bias. This provides more information and helps to increase the sensitivity of a surveillance system.

In this study, a spatial zero-inflated Poisson model (SZIP) was adjusted to the temporal patterns of reports on ASF-positive wild boar carcass in the Republic of Korea. The study aimed to (1) disentangle the factors affecting the presence and reporting of ASF-positive carcasses, (2) estimate the prevalence and sensitivity of the surveillance system, and (3) understand the dynamics of ASF infection among the wild boar population.

## MATERIALS AND METHODS

The reported data for wild boars were retrieved from the surveillance system database developed by the Ministry of Environment, Republic of Korea. The collected data included the type of report (public reporting or active searching), report date, diagnosis date, species (e.g., wild boar), type of specimen (carcass of or captured wild boar), administrative addresses, and geographic coordinates. Among the ASF-positive cases reported, only the ASF-positive cases identified from carcasses were extracted for analysis. Between the first case of ASF in wild boar carcasses reported on October 02, 2019, and April 24, 2020, a total of 569 cases of infected carcasses were reported.

To divide the study period according to the reporting patterns of ASF-positive carcasses, test for structural break was conducted for the time series of ASF-positive carcasses. The structural breaks are the time points when the time series data changes abruptly (Zeileis et al., 2003). These can be identified by minimizing the residuals of the sum of squares of the single or multiple linear regressions for the time series. Considering the residuals of the sum of squares and the complexity of the models, the number and time points of structural breaks were selected based on the Bayesian Information Criteria (BIC). The test was conducted with the *“strucchange”* package (Zeileis et al., 2001) in R software version 4.0.2 (R Core Team, 2020).

Considering the imperfect detection in the surveillance of wild boars, the regions that reported ASF-positive carcasses and their neighboring regions were selected as the study sites and were marked using the 2^nd^ level administrative regional (si-gun-gu) boundaries. These regional boundaries followed the 2019 Administrative District Boundary for the Republic of Korea from Statistics Korea (Statistics Korea, 2019). Since the movement of wild boars does not follow the boundary of the administrative regions and low spatial resolution could limit the spatial analyses, the study region was partitioned into 1,237 regular hexagons with a 3-km diameter. For each of the two study periods, defined by the structural break analysis, the number of the ASF-positive carcasses that were reported was counted in each hexagon.

Zero-inflated Poisson (ZIP) regression model is a mixture model of a logistic and a Poisson regression model. Its associated probability distribution of the counts can be expressed as follows:

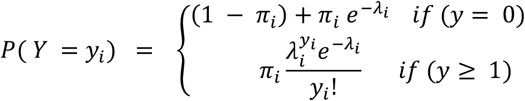

where, *i* is the index for the hexagon; *y_i_* is the observed number of the ASF-positive carcasses in the hexagon *i*; *π_i_*, is the probability of presence of ASF-positive wild boar carcasses in a hexagon *i*; *λ_i_* is the average number of reports of ASF-positive carcasses for a hexagon *i* given the presence of at least one the ASF-positive carcass. The model can be extended to account for covariates in the logistic and Poisson parts as follows:

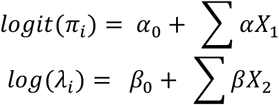

where, *α*_0_ and *β*_0_ are intercepts, *α* and *β* are vectors of coefficients, and *X*_1_ and *X*_2_ are vectors of covariates for logistic and Poisson parts, respectively. Note that *X*_1_ and *X*_2_ can be different sets of covariates, which can help to disentangle the factors for the presence and reports of ASF-positive carcasses. The model divides the zeros into two categories: true zero and false zero. The former comes from the true absence of ASF-positive carcasses, leading to no reports of case (i.e., true negative). The latter comes from affected hexagons (i.e. where ASF is present) where no ASF-positive carcasses have been found. (i.e., false negative). The set of covariates involved in the Poisson part (*X*_2_) allows *λ* (the average number of reports of ASF-positive carcasses) to vary between hexagons and therefore account for both heterogeneous abundance of positive carcasses and for heterogeneous surveillance effectiveness. Note that amongst the variables in *X*_2_, the factors associated with the abundance of positive carcasses cannot be distinguished from the those associated with surveillance effectiveness. Similar to (Vergne et al., 2014; Vergne et al., 2016), we assumed a spatial autocorrelation on the probability of presence of ASF-positive carcasses. Therefore, we included an intrinsic conditional autoregressive component in the logistic part of the ZIP, such as the following:

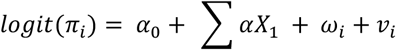

where, *ω_i_*, is a spatially structured random effect and *v_i_*, is an unstructured random effect. For the former, the spatial adjacency structure was assumed based on the first-order contiguity between the hexagons (Besag et al., 1991). The latter was specified based on Gaussian distribution with a mean of zero.

The sensitivity of the surveillance system for each hexagon 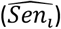, which is defined as the detection probability of at least ASF-positive carcasses in affected hexagons, can be estimated for each hexagon as one minus the probability of no report when ASF-positive carcasses are present 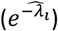, as in 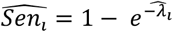. Therefore, the probability of being a false negative can be expressed as follows: 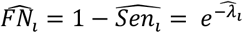. Note that specificity is assumed to be perfect in the zero-inflated model; thus, all ASFV-positive wild boar carcasses are true positives. The probability that at least one case was reported in each hexagon can be estimated with the risk multiplied by the sensitivity of the surveillance system 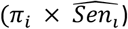. The hexagon-level prevalence 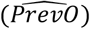 was estimated with the following formula (Vergne et al., 2014):

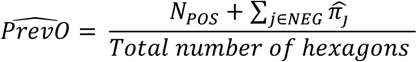

where, *NEG* correspond to hexagons with no reported cases, 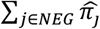 is estimated number of affected hexagons with no reported cases (i.e., the number of false negative hexagons); *N_POS_* is the number of affected hexagons with reports of the ASF-positive carcasses. The hexagon-level sensitivity of the surveillance system 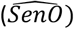 as follows:

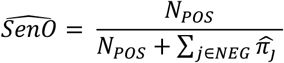

### Covariates

Considering the symptoms of ASF-infected wild boars, such as fever, dehydration, and hemorrhage, the behaviors of infected wild boars would be associated with those of the feverish animals preference for a cool, humid, and aqueous environment (Morelle et al., 2019).). Thus, the covariates were selected based on these characteristics and previous studies (Morelle et al., 2019; Podgórski et al., 2019; Cukor et al., 2020).

The habitat suitability for wild boars was retrieved from the species distribution model to identify the distribution of wild boars (Eu-Tteum and Son-Il, 2020). The model provides values with a 1-km-resolution in a raster format, and the values summed at each hexagon were utilized in the analyses. This variable was included in both parts of SZIP, regardless of statistical significance, to adjust for the confounding effect of wild boar distribution on the associations between risk factors and the probability of presence and the number of reports. Moreover, elevation and slope data were analyzed (Morelle et al., 2019). The elevation data were obtained from the Space Shuttle Radar Topography Mission 1 Arc-Second Global data version 3.0 (JPL, 2013). These data provide the elevation value with 30-m-resolution raster data. The slope was calculated with the same resolution as that of the data. To estimate vegetation distribution and its intensity (Podgórski et al., 2019; Cukor et al., 2020), Enhanced Vegetation Index (EVI) was obtained from satellite products of the Moderate Resolution Imaging Spectroradiometer (MODIS), NASA. EVI representing the level of vegetation activity was obtained from the Terra MODIS MOD13Q1 Version 6 product with a geographic resolution of 250 m (Didan, 2015).

Human density could be associated with the distribution of wild boars or the report of ASF-infected carcasses (Podgórski et al., 2019). These data were retrieved from the Gridded Population of the World Version 4, Center for International Earth Science Information Network, Columbia University (Center for International Earth Science Information Network - CIESIN - Columbia University, 2018) with a geographic resolution of 30 arc-second (approximately 1 km). Polygons for the rice paddy, surface water, and wetland in the study region were obtained from the Ministry of Environment and utilized to calculate the areas distributed in each hexagon. The covariates could be related to the distribution of wild boars and behaviors of ASF-infected wild boars (Morelle et al., 2019; Cukor et al., 2020).

The minimal distance from North Korea to the centroid of the hexagons was included. Using the shape file of North Korea downloaded from Global Administrative Areas version 3.6 (Global Administrative Areas, 2012), which represented the administrative map in 2018, the minimal distance from each centroid of a hexagon to North Korea was calculated. The heat load index (HLI) is defined as the estimate of potential annual direct incident solar radiation. To assess the temperature at deathbed (Cukor et al., 2020), the HLI considers slope, and aspect, ranging from 0 (coolest) to 1 (hottest) (McCune and Keon, 2002). Land surface temperature at day (LSTD) and night (LSTN) were retrieved from the Terra MODIS MOD11A1 product Version 6 (Wan et al., 2015). This product provides the raster format with a 1 km resolution. To analyze the relationship between the deathbed and humidity, precipitation, and Normalized Difference Water Index (NDWI) were retrieved. Precipitation data were retrieved from the Automatic Synoptic Observation System, Korea Meteorological Administration (Korea Meteorological Administration, 2020). As the data consisted of observed values with their geographic coordinates, the spatial, continuous surfaces of the precipitation in the study regions were estimated using ordinary kriging. NDWI was calculated, which indicated the water content of vegetation, using near infrared (858 nm) and shortwave infrared (1650 nm). Each of these wavelengths corresponded to bands 2 and 6, respectively, in the Terra MODIS MOD09A1 Version 6 product, with a 500-m resolution (Boschetti et al., 2014; Vermote, 2015)

The median of distributions of time-varying variables, such as EVI, LSTN, LSTD, precipitation, and NDWI, were extracted from each source at the highest temporal resolution for the total study period. Consistent variables, including habitat suitability for wild boars, elevation, slope, aspect, area of rice paddy, surface water and wetland, minimal distance to North Korea, HLI, and human density were extracted from each source and equally analyzed for both the study periods. All values of the variables, spatially corresponding to a hexagon, were extracted, and the median of the values was assigned for the hexagons. To account for non-linear association, the values were categorized into three levels with a similar number of hexagons at each level. The area of the wetland was categorized into two levels (none or present) because 834 of the total hexagons had no wetlands.

### Model building

The models were built using the following: univariable analysis, collinearity analysis, and multivariable analysis, with a forward selection procedure. Collinearity between the variables was assessed using the Kendall rank correlation test. One of the pair of variables with a correlation of over 0.7 was included in the univariable analysis. In the univariable analysis, all variables were tested independently using the Poisson and the logistic parts of the ZIP regression model. In the univariable analysis for both time periods, besides habitat suitability of wild boar, the spatial and non-spatial random terms were included in the logistic part of the regression analysis. In particular, for univariable analysis of the second study period, the spatiotemporal and temporal autocorrelations were adjusted; this was achieved using the binary variable, indicating whether ASF-positive carcasses were reported in the first study period in the hexagon and their neighborhoods (ST_variable), and the continuous variable for the number of reports during the first study period in the hexagon (T_report) as a confounder in the logistic and Poisson parts, respectively. Variables associated with at least one coefficient for which zero was not included in the 80% credible interval (CrI) of its posterior distribution were included in the multivariable analysis. In multivariable analysis, a forward selection procedure was conducted to build the final models based on the Deviance Information Criteria (DIC), which quantified the balance between the complexity and the fit of the model (Spiegelhalter et al., 2002). The best model was the one with DIC less than two points greater than that of the model with the smallest DIC. Similar to the univariable analysis, to adjust confounding effects, habitat suitability for wild boars was included in the Poisson and logistic parts of the regression analysis.

All analyses were performed with Bayesian Markov Chain Monte Carlo simulation (MCMC). The MCMC was performed in R software with the *“R2OpenBUGS”* package (Sturtz et al., 2010). The prior distributions for the coefficients of the SZIP were set as non-informative normal distributions with a mean of 0 and a variance of 10. For spatial and non-spatial random effects, the prior distribution was assumed to follow the inverse Gamma distribution with a mean of 10 and a variance of 100.

Three chains of 100,000 iterations were sampled with 20,000 burn-in. Convergence was assessed by checking the trace plot visually and using the Gelman-Rubin-Brooks diagnostic (Gelman et al., 2013). The posterior distribution of the parameters is summarized with the median and the 95% CrI values. Furthermore, for each hexagon, median values of the risk, sensitivity, the probability that at least one case was reported, and the probability of false negative report were plotted as choropleth maps. The discriminatory performance of the model was assessed using the area under the curve (AUC) of the receiver operating characteristic curves for the estimated probability that at least one case was reported in the hexagons. To compare the hexagon-level prevalence of ASF-positive wild boar carcasses, the hexagon-level sensitivity of the surveillance system and the hexagon-level proportion of false negative reports between the two study periods were plotted against each of the posterior distributions. Scatterplot for the probability of the presence of ASF-positive wild boar carcasses between the first and second periods was showed.

Because of substantial computational demand, the simulations were run with parallel computing using *“parallel”* and *“dclone”* (Solymos et al., 2019) packages in R software.

## RESULTS

The results of the structural break analyses are shown in Figure 1. Only one date—January 19, 2020—was identified as a structural break based on the BIC. The study periods were divided into two parts using this break: the first period was from October 02, 2019, to January 18, 2020, and the second period was from January 19, 2020, to April 28, 2020. During the first period, the total number of reports of ASF-positive carcasses was 85, distributed among 37 hexagons. For the second period, 484 of the ASF-positive carcasses were reported in 80 hexagons.

**Figure 1.**
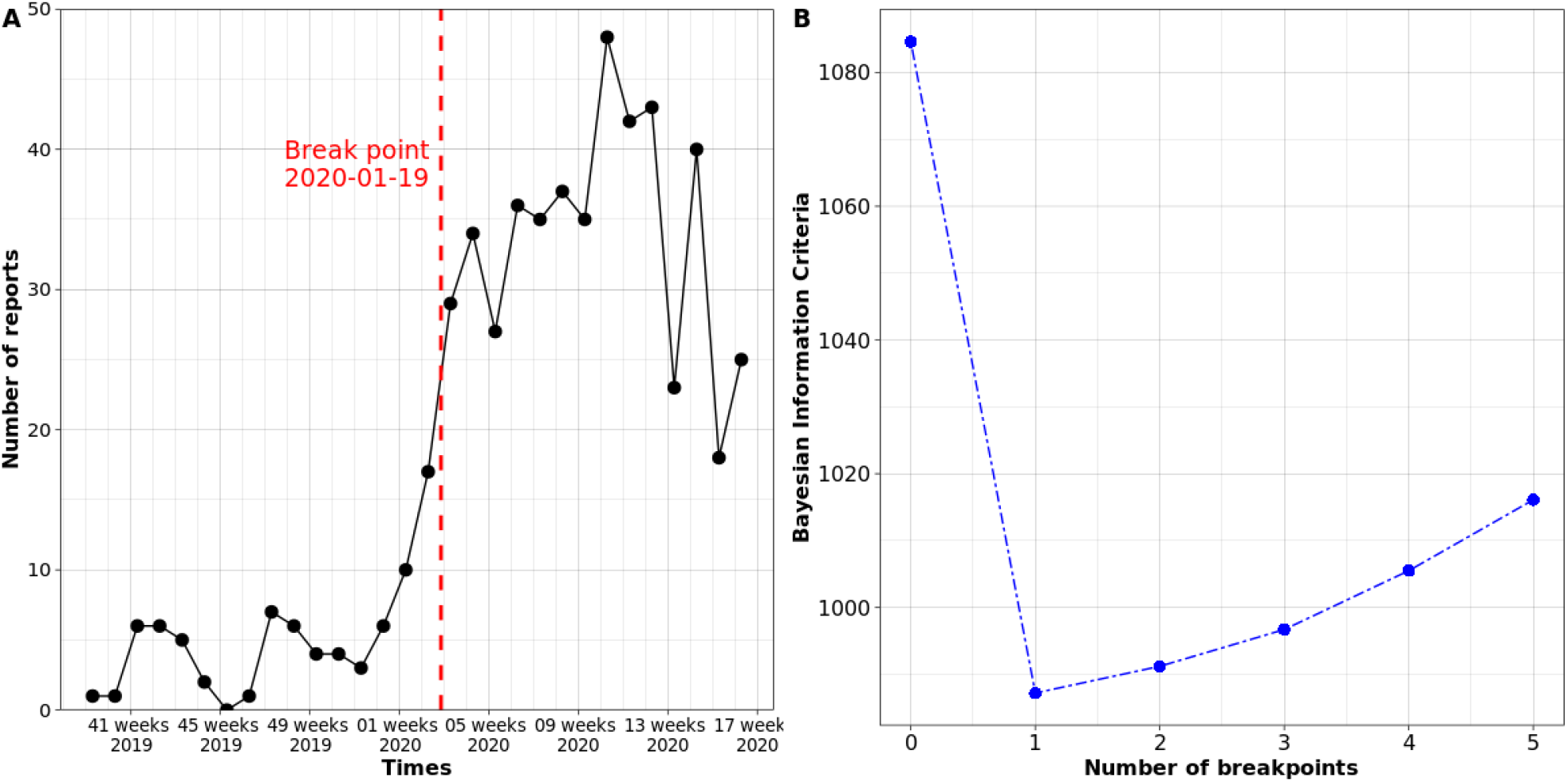
Time series plot for the number of reports per week from October 2019 to April 2020 (A): Red dashed vertical line shows the break point on 19^th^ January 2020. Bayesian information criteria estimated with each number of breakpoints (B).

The results of univariable analyses for the first and second periods are presented in Table S1 and S2: for the first period, elevation, slope, area of rice paddy and surface water, minimal distance to North Korea, HLI, human population density, EVI, LSTD, precipitation, and NDWI were significant at 20% level in both the Poisson and logistic parts of the regression analysis; elevation, slope, area of rice paddy, presence of wetland, minimal distance to North Korea, HLI, human population density, EVI, and precipitation were significant at 20% level only in the Poisson part of the analysis; area of rice paddy, minimal distance to North Korea, human population density, LSTD, LSTN, precipitation, and NDWI were significant at 20% level in only the logistic part of the analysis. For the second study period, elevation, area of rice paddy and surface water, presence of wetland, minimal distance to North Korea, HLI, human population density, EVI, LSTD, LSTN, precipitation, and NDWI were significant at 20% level in both the Poisson and logistic parts of the regression analysis; elevation, area of rice paddy and surface water, minimal distance to North Korea, HLI, human population density, EVI, LSTD, precipitation, and NDWI were significant at 20% level in only the Poisson part of the analysis; presence of wetland, minimal distance to North Korea, human population density, LSTD, and NDWI were significant at 20% level in only the logistic part of the analysis. Among the significant variables at 20% in the univariable analyses (Table S3), LSTD was highly correlated with elevation and slope for the first study period (Table S4); for the second study period, LSTD was highly correlated with slope. LSTD was excluded in the multivariable analyses.

The study region was located in the northern regions of South Korea, bordering North Korea (Figure 2). The results of the SZIP for the first study period are presented in Table 1. For the first period, only the distance from North Korea to the centroid of hexagons was identified as a significant factor for the presence of the ASF-positive wild boar carcasses. The estimated odds ratios (OR) were 0.12 (95% CrI: 0.01–0.95) and 0.01 (95% CrI: 0.00–0.30) for the hexagons located between 14.62 km and 28.16 km, and those located between 28.16 km and 76.77 km, respectively, relative to the estimated OR of hexagons located between 0 to 14.62 km from North Korea. However, habitat suitability for wild boar showed negative association without any statistical significance, with an OR of 0.21 (95% CrI: 0.02–1.45) and 0.07 (95% CrI: 0.00–1.33), compared to the first level of habitat suitability for wild boar. In the Poisson part of the regression analysis, the habitat suitability for wild boar, HLI, and human population density were identified as statistically significant risk factors for the number of reported cases of ASF-positive carcasses in the affected hexagons. The estimated incidence rate ratio in the hexagons with the second- and third-level of habitat suitability for wild boar were 2.33 (95% CrI: 1.02–4.85) and 8.75 (95% CrI: 1.44– 55.54), respectively, compared to the first level of habitat suitability for wild boar. For HLI, only the third level of HLI showed a statistically significant positive association (IRR: 0.40, 95% CrI: 0.17–0.93). The second level of HLI showed a positive association but was not significant (IRR: 1.50, 95% CrI: 0.80–2.98). Although human population density showed a positive association, only the density of 61.26–9448.68 person per km^2^ was statistically significant (human density of 24.77–61.26 person per km^2^, IRR: 3.23, 95% CrI: 0.75–17.00; human density of 61.26–9,448.68 person per km^2^, IRR: 4.99, 95% CrI: 1.09–27.88).

**Figure 2.**
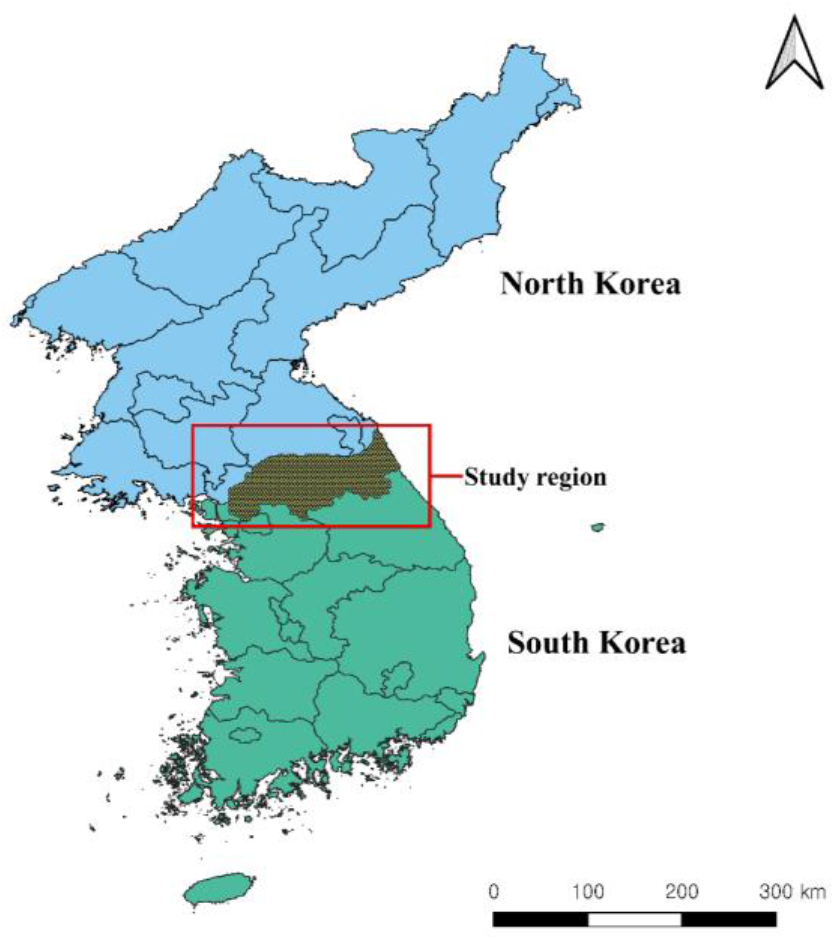
Map showing the location of study region in South and North Korea.

**Table 1.** Results of zero-inflated Poisson model for the first period

| Variables | Category | Number of hexagons | Median of odds ratio | 95% CrI* | Median of incidence rate ratio | 95% CrI |
| --- | --- | --- | --- | --- | --- | --- |
| Suitability of wild boar | [0.29 - 0.46) | 408 |  | Ref. |  | Ref. |
|  | [0.46 - 0.59) | 408 | 0.21 | 0.02 – 1.45 | 2.33 | 1.02 – 4.85 |
|  | [0.59 - 0.85] | 421 | 0.07 | 0.00 – 1.33 | 8.75 | 1.52 – 56.43 |
| Distance to North Korea (km) | [0 – 14.62) | 408 |  | Ref. |  |  |
|  | [14.62 – 28.16) | 408 | 0.12 | 0.01 – 0.95 |  | - |
|  | [28.16 – 76.77] | 421 | 0.01 | 0.00 – 0.30 |  |  |
| Heat load index | [0.62 – 0.81) | 408 |  |  |  | Ref. |
|  | [0.81 – 0.84) | 408 | - |  | 1.50 | 0.80 – 2.98 |
|  | [0.84 – 0.99] | 421 |  |  | 0.40 | 0.17 – 0.93 |
| Human population density (person per km<sup>2</sup>) | [15.68 – 24.77) | 408 |  |  |  | Ref. |
|  | [24.77 – 72.36) | 408 | - |  | 3.23 | 0.75 – 17.00 |
|  | [72.36 – 9448.68] | 421 |  |  | 4.99 | 1.09 – 27.88 |
| **Spatial component** | Variance | 95% CrI |  |  |  |  |
| Spatial random term | 12.99 | 2.71 – 51.14 |  |  |  |  |
| Non-spatial random term | 0.19 | 0.03 – 2.93 |  |  |  |  |

\*CrI: Credible interval

Table 2 shows the results of the SZIP for the second study period. In the logistic part of the analysis, only ST_variable, representing whether ASF-positive carcasses were reported in a hexagon and its neighborhoods during the first study period, showed a significant positive association with an OR of 29.90 (95% CrI: 5.92–220.96). Habitat suitability for wild boar showed a negative association with the risk of ASF; however, the association was not statistically significant (second level of habitat suitability for wild boar: OR: 0.75, 95% CrI: 0.08–5.11; third level of habitat suitability for wild boar: OR: 0.16, 95% CrI: 0.01–2.72). T_report showing the number of reports of ASF-positive carcasses in a hexagon during the first period, habitat suitability for wild boar, elevation, and HLI were identified as statistically significant factors associated with the number of reported carcasses in the affected hexagons. The estimated IRR of T_report was 1.13 (95% CrI: 1.07–1.20). Habitat suitability for wild boar showed a weak positive association (second level of habitat suitability for wild boar: OR: 2.13, 95% CrI: 1.58–2.94; third level of habitat suitability for wild boar: OR: 1.26, 95% CrI: 0.72–2.13). HLI showed a significant positive association (second level HLI: IRR: 2.92, 95% CrI: 2.03–4.28; third level HLI: IRR: 3.00, 95% CrI: 2.08–4.44). Although elevation showed a negative association, only the third level of the elevation was significantly significant (second level elevation: IRR: 0.88, 95% CrI: 0.63–1.24; third level elevation: IRR: 0.63, 95% CrI: 0.41–0.96).

**Table 2.**
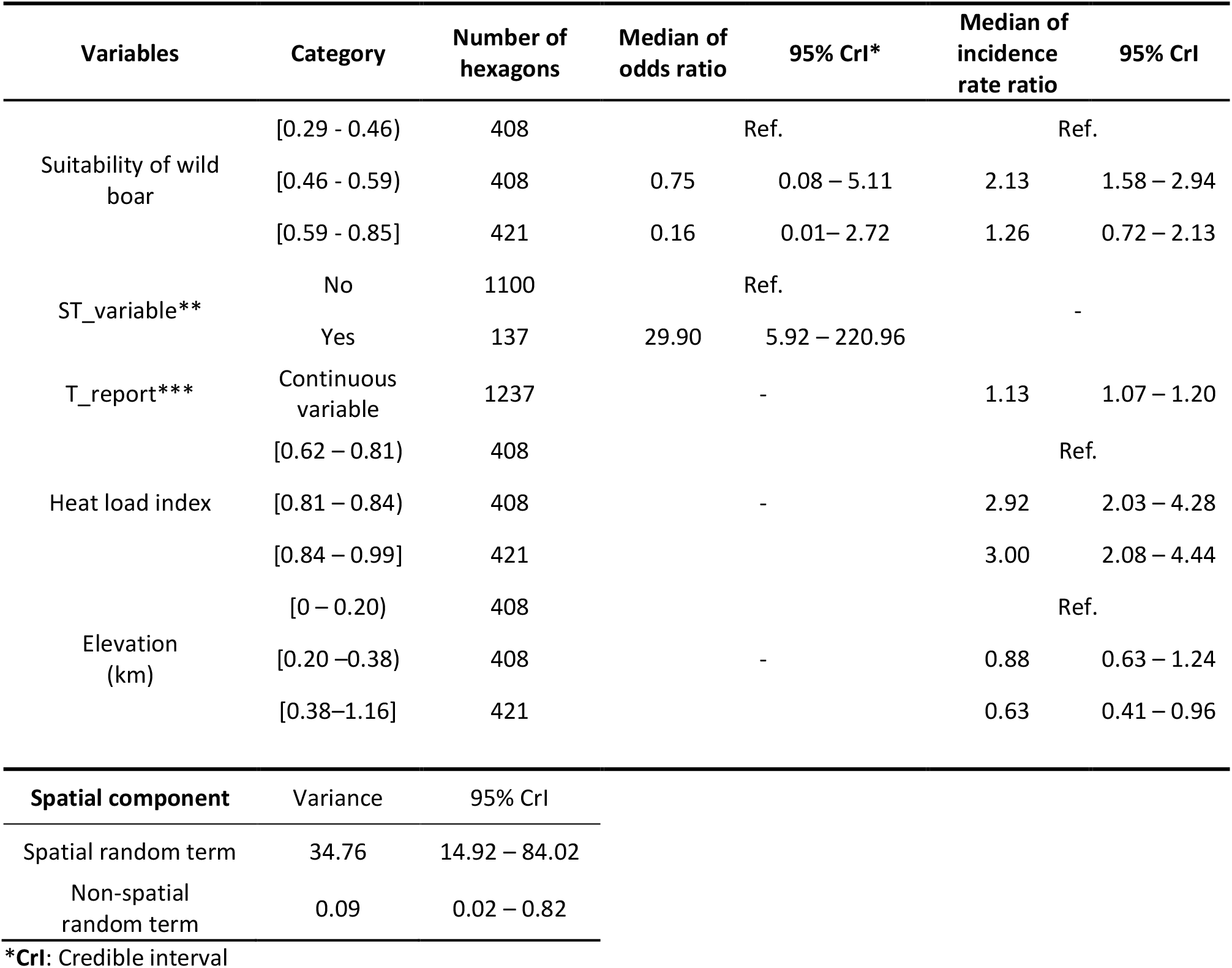
Results of zero-inflated Poisson model for the second period

The AUCs of the models were 0.98 (95% CrI: 0.96–0.99) and 0.98 (95% CrI: 0.97–0.99) for the first and second study periods, respectively. With the estimated models, the median values of the risk, sensitivity, and the probability that at least one case was reported were plotted with the number of reports (Figure 3). For the first study period, the hotspots with ASF-positive carcasses were located in the north-west and central north regions (Figure 3E). Sensitivity was lower in the central regions than that in the other regions. Moreover, the regions surrounding those with reported cases had a low surveillance sensitivity (Figure 3G). During the second study period, the maximum value of the probability of the presence of ASF-positive wild boar carcasses in the hexagons was higher than that during the first study period. The hotspot of the central north regions had moved to the south-east (Figure 3F). The overall sensitivity was higher in the second study period than that during the first study period. However, hexagons with relatively low sensitivity were present in the eastern regions (Figure 3H). Figure 4 shows the probability of false negative in each hexagon during the first and second study periods. During the first study period, there were hexagons with a high probability of false negatives in the central and northwest regions where ASF-positive carcasses were reported. Overall, the probabilities of false negative during the second period were lower than that during the first period. Nonetheless, there were hexagons with a relatively high probability of false negatives in the central and northeast regions where ASF-positive carcasses were reported.

**Figure 3.**
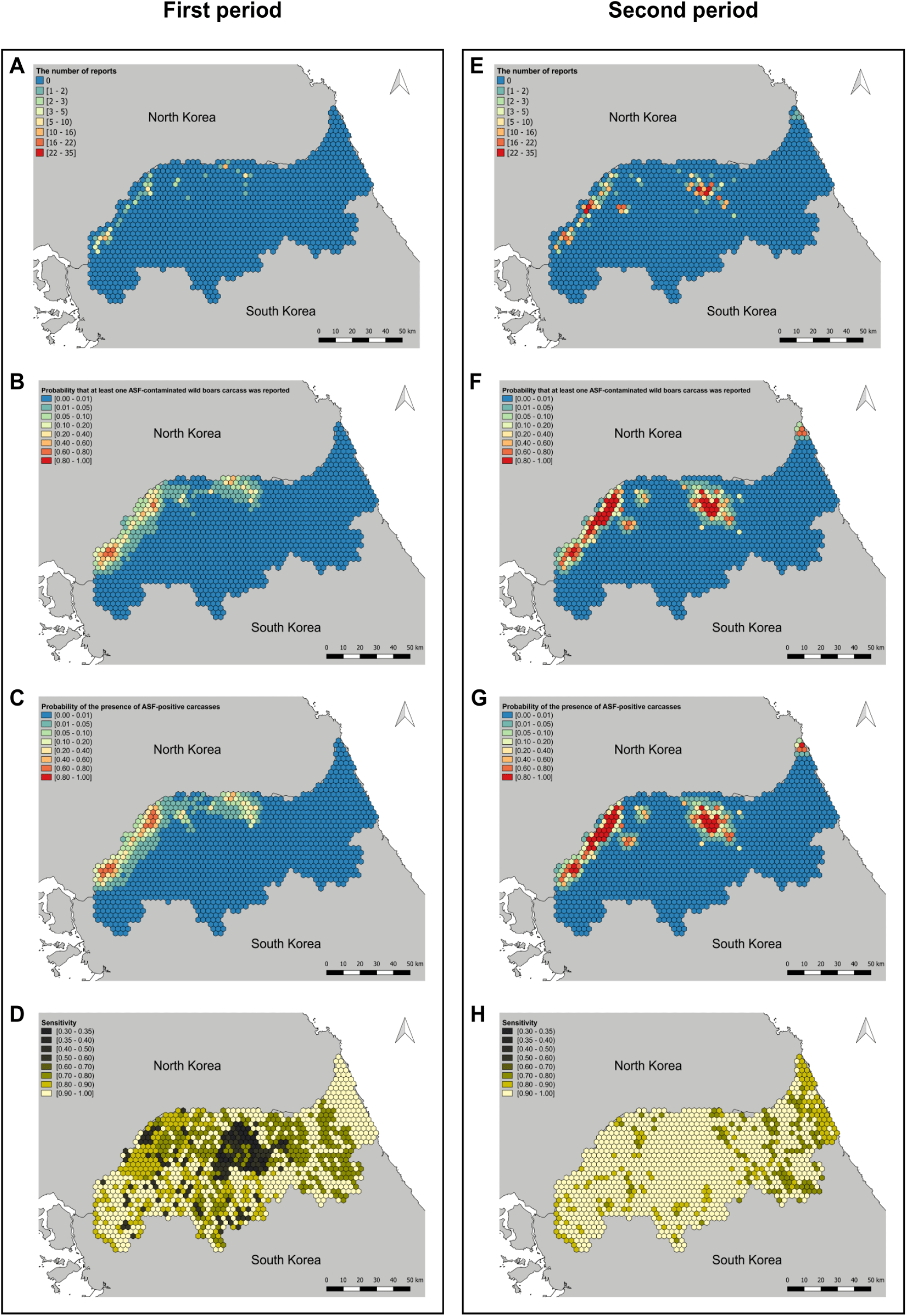
Choropleth maps of the number of reports of African swine fever (ASF) – positive wild boar carcasses (A, E for each period),, the median value of the probability of at least on report of ASF-positive wild boar carcasses (B, F for each period), the median value of the probability of presence of ASF-positive wild boar carcasses (C, G for each period), and the median value of the sensitivity (E,H for each period) for the first and second periods.

**Figure 4.**
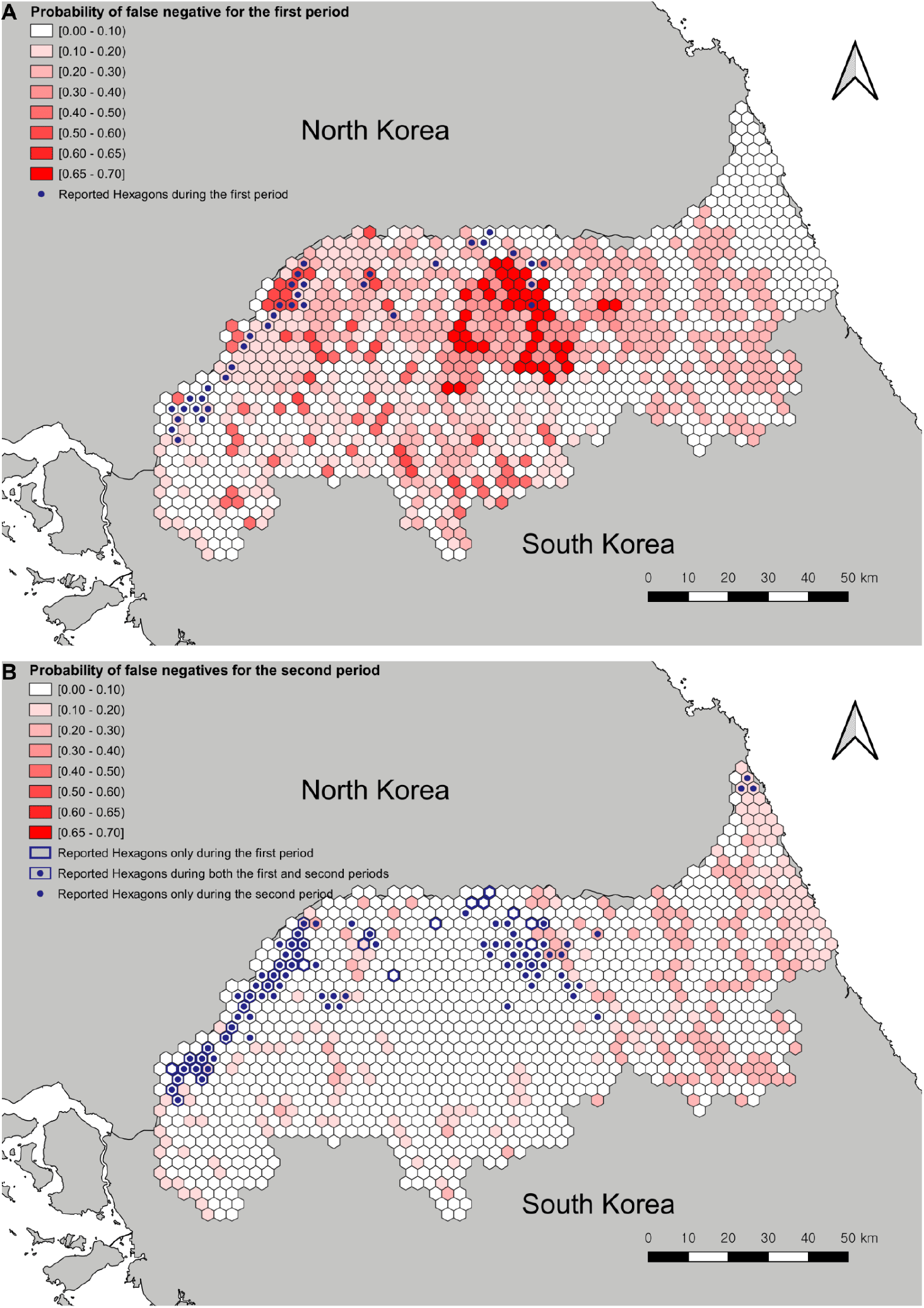
Choropleth map of the probability of false negative during the first (A) and second period (B)

The hexagon-level prevalence, sensitivity, and proportion of false negative for each period are presented in Table 3 and Figure 5. The number of affected hexagons with no carcasses reported was not statistically different between the two study periods: first period, 39 (95% CrI: 29–53); and second period, 30 (95% CrI: 20–43). The hexagon-level prevalence of ASF-positive carcasses was significantly greater during the second period than that during the first (0.09, 95% CrI: 0.08–0.10 vs 0.06, 95% CrI: 0.05–0.07). Hexagon-level sensitivity increased from 0.49 (95% CrI: 0.41–0.57) in the first period to 0.73 (95% CrI: 0.66–0.81) in the second period. The proportion of false negative was not statistically different between the first (0.03, 95% CrI: 0.02–0.04) and second study periods (0.02, 95% CrI: 0.02–0.03). Figure 6 shows a scatterplot representing a positive linear relationship between the probabilities of the presence of ASF-positive carcasses in the first and second study periods.

**Figure 5.**
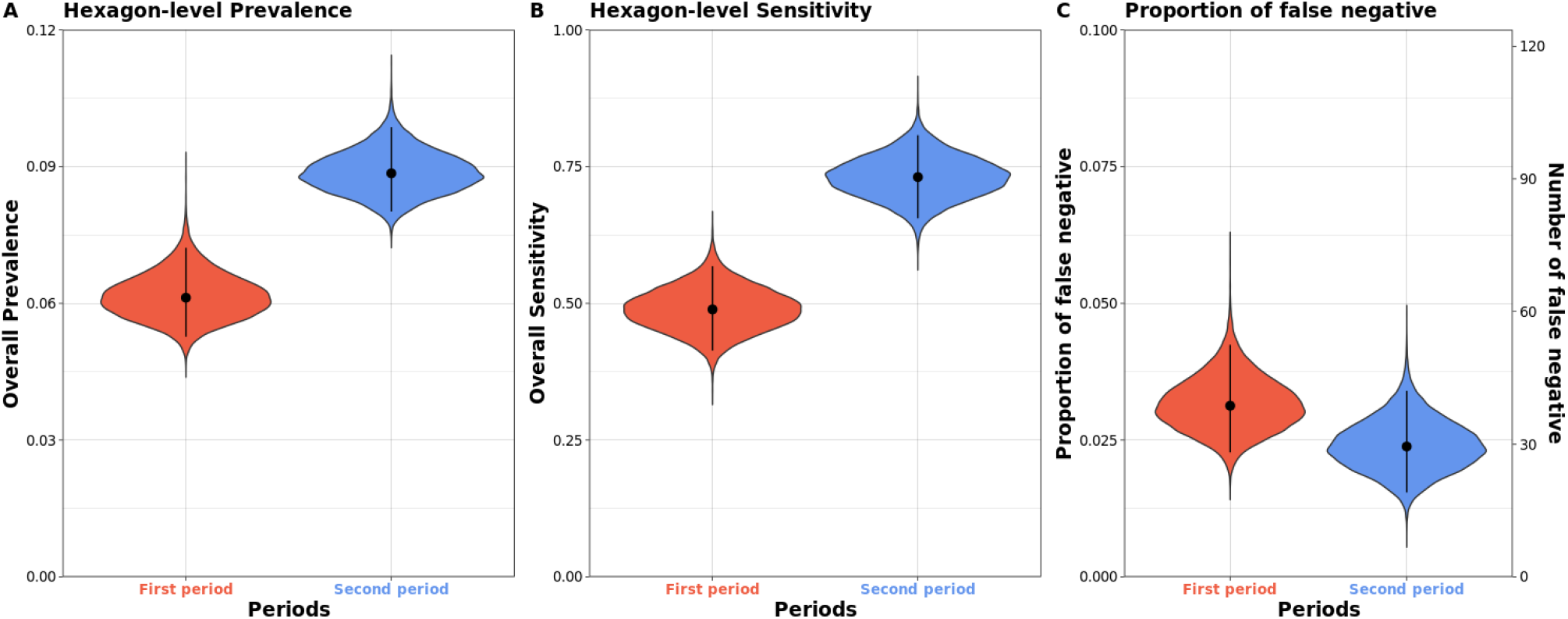
Violin plots for the posterior distribution of the hexagon-level prevalence (B), hexagon-level sensitivity (A), and proportion and number of the affected but undetected hexagons (C) for the first and second periods.

**Figure 6.**
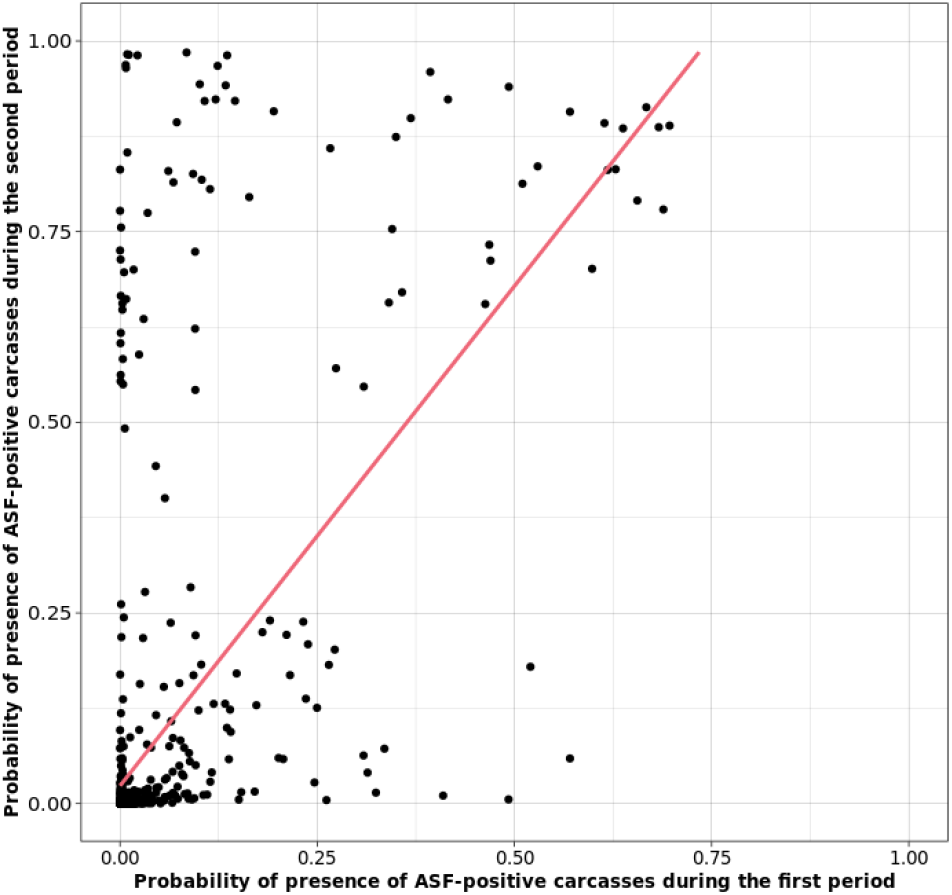
Scatter plot for the relationship between the probability of presence of ASF-positive wild boar carcasses for the first and second periods.

**Table 3.** The hexagon-level prevalence, sensitivity, and proportion of false negative for each period.

| Value | The first period<br>(95% CrI*) | The second period<br>(95% CrI) |
| --- | --- | --- |
| Total number of hexagons | 1,237 | 1,237 |
| Number of reported hexagons<br>(True positive) | 37 | 80 |
| Number of undetected hexagons<br>(False negative) | 39 (29 – 53) | 30 (20 – 43) |
| Hexagon-level prevalence | 0.06 (0.05 – 0.07) | 0.09 (0.08 – 0.10) |
| Hexagon-level sensitivity | 0.49 (0.41 – 0.57) | 0.73 (0.66 – 0.81) |
| Proportion of false negative | 0.03 (0.02 – 0.04) | 0.02 (0.02 – 0.03) |

\*CrI: Credible interval

## DISCUSSION

To control ASF dynamics among wild boar populations, the Korean government has developed and operates the surveillance systems. However, wildlife surveillance has limitations of imperfect disease detection because of the dependence on public reporting and limited resources; thus, it is highly likely to create confusions as to whether the identified risk factors are associated with the risk or reporting of ASF-positive carcasses (Combelles et al., 2019), which can also result in uncertainties about the dynamics of ASF among wild boar populations. In this study, we analyzed the spatial distributions of deathbed of ASF-infected wild boars for successive periods, while adjusting for the imperfect detection: (1) risk factor associated with the presence of ASF-positive carcasses was changed from proximity to North Korea to whether ASF-positive carcasses were reported in a hexagon and its neighborhoods during the previous study period; (2) habitat suitability for wild boar, human population density, HLI, and elevation were associated with the number of reports of ASF-positive carcasses in the affected hexagons; (3) hexagon-level prevalence of ASF-positive carcasses increased and the hexagon-level sensitivity also improved during the study period.

For the first study period, our results emphasized that the infection risk of ASFV is significantly increased with decreasing distance from the North Korean border, suggesting that the ASFV could have been introduced from North Korea. This is consistent with the fact that North Korea reported an ASF outbreak on May 30, 2019, before the ASF epidemic in South Korea (World Organisaztion for Animal Health (OIE), 2019). South Korea is geographically connected to Asia through North Korea. Although the buffer zone between the two countries, which is also called the Demilitarized Zone, can be a physical barrier to the movement of ASF-infected wild boars, there remains a risk of introduction of ASFV through scavengers or other mechanical routes.

As expected, the results also showed that the habitat suitability for wild boar is associated with the number of reports of ASF-positive carcasses in the affected hexagons. This is likely due to the higher contact rates between wild boar resulting in a higher incidence and therefore a higher number of cases, leading to an increased number of reports of ASF-positive carcasses. Moreover, more ASF-positive carcasses were reported in the affected hexagons with low HLI. This is likely because ASF-infected wild boars prefer high-moisture and cool-temperature environments because of the clinical symptoms of high fever and dehydration (Gabriel et al., 2011), leading to an increased abundance of infected wild boars and therefore ASF-positive carcasses, which is consistent with previous studies (Podgórski et al., 2019; Cukor et al., 2020). Furthermore, this observation might be due to the fact that ASFV is more resistant to colder environments, which means that the virus remains active for a longer time, generating more opportunities for transmission (Bellini et al., 2016). Moreover, it takes more time for wild boar carcasses to decompose in shade than in sunlight (Probst et al., 2020), which can make detection easier and therefore increase the number of reported carcasses. Usually, human population density is a proxy for human activities and therefore passive disease detection in veterinary epidemiological research (Ward et al., 2006; Gilbert et al., 2008; Martin et al., 2011; Dhingra et al., 2016). Consistently, we found that human population density was associated with the number of reported ASF-positive carcasses in the affected hexagons. During the first study period, the surveillance was highly dependent on the passive disease detection (i.e., public reporting). Moreover, the reporters and hunters employed by the government were likely to get more help from local residents to locate carcasses in more densely populated regions.

During the second study period, the fact that the carcasses were already reported during the first period in a hexagon or its neighboring hexagons was a strong risk factor for the presence of ASF-positive carcasses. Unlike during the first study period, the distance to North Korea was not associated with the presence of ASF-positive wild boar carcasses, suggesting that disease distribution was driven by the previous phase of the epidemic rather than by potential re-introductions of ASFV from North Korea.

The results of the second period also showed that in the affected hexagon, the greater the number of reports during the first period, the greater the number of reports during the second period. This could be due to the enhanced surveillance that was focused on previously affected regions. Moreover, HLI was found to be positively associated with the number of reported carcasses in the affected hexagons, while no such observation was noted during the first study period. This could be due to seasonal effects of HLI. Unlike during the first period, the second period included spring season. As the regions with high HLI receive more solar radiation, they would have more dense vegetation than the other regions (Szeicz, 1974; He et al., 2017), and consequently have moist and cooler microclimate (Aussenac, 2000). Thus, these environments can attract more ASF-infected wild boars. Alternatively, it can be interpreted that the infected wild boars prefer this type of environment to hide from the hunters or predators. Thus, more ASF-positive carcasses might be reported in such regions. The results also suggest that among the affected hexagons, the higher the altitude of a region, the lower the number of reports. As surveillance was still highly dependent on human resources, it may be difficult to report the carcasses in regions that are difficult to access for people. Furthermore, habitat suitability for wild boar showed a weak statistical association with the number of reports. This may be due to the enhanced surveillance for wild boars during the second study period, which induces changes in the population behaviors. Wild boars flexibly adjust their activity and home ranges to human activities, such as hunting and recreation, to protect themselves from risk (Scillitani et al., 2009; Thurfjell et al., 2013; Johann et al., 2020). Owing to intensive hunting condition, wild boars could have left their original home-range and shifted to other areas where they could hide from the hunters (Thurfjell et al., 2013). This makes the spatial distribution of wild boars unstable and interspersed (Scillitani et al., 2009).

The results also suggested that the increased number of reported ASF-positive carcasses from the first to the second study periods were the combined results of the increased prevalence of ASF and an improved sensitivity of the surveillance system. Taken together with the maps for the probability of presence of ASF-positive carcasses and its relationship between the first and second study periods, the probability of false negative reports, and the increased prevalence for the both periods, it can be stated that ASF infection had spatially spread centered on the previously affected regions, especially in the southward direction. Although it is favorable that the overall sensitivity of the surveillance system increased in the second period, it should be noted that some regions remained undetected. These might have contributed to the spatial spread. The movement of the hotspot towards the south-eastern direction in the second study period may be explained by the following possibility: there were hexagons with low surveillance sensitivity around the central north hotspot in the first period. Thus, ASFV could have been introduced into these regions and remained undetected during the first study period. The undetected ASF-positive carcasses could have contributed to the infectious events in these regions, which could have generated the hotspot identified in the second study period. Similarly, the undetected hexagons in the second period might contribute to spatial spread in the future: the carcasses undetected in the northeast regions in the second period could contribute to the spread of ASFV in the southern direction because of low surveillance sensitivity in these regions. Furthermore, the “Taebaek mountain ranges” are present around these regions, starting from North Korea to the middle of South Korea. Since ASFV infection among wild boars could follow the boundary of these mountain ranges (NIBR, 2017), controlling disease spread may be more demanding due to the high elevation and low human population density. Considering that the study regions include the border with North Korea, there are areas with unique characteristics: Demilitarized Zone (2 km from the border) and Civilian Control Zone (10 km from the Demilitarized Zone), which correspond to the reference level of variable of the distance to North Korea. In these regions, there were no residents because of military reasons. Moreover, there are landmines in these regions. Thus, even if the probability of the presence of the ASF-positive carcasses was high, it would have been difficult to search for such carcasses. Therefore, during the first period, it can be assumed that there were a significant number of undetected cases in these regions. These undetected carcasses could be a driver of the endemicity of ASF infection among wild boars. This study had two main limitations. First, because of military restrictions, many types of spatial data were unavailable for distance below 25 km from the North Korean border, which covers 54% of the study region. Consequently, many variables that are potentially related to observed distribution of ASF-positive carcasses, such as road length, density of domestic pig farms, and landmine distribution could not be included. Thus, most of the variables in this study were dependent on remote sensing data. Second, we did not account for the death time point of the carcasses in the analyses. This might result in bias in the results since carcasses found during the second study period could be due to the death of a wild boar during the first period. However, the data of time of death was not available as this estimation is incredibly challenging owing to its dependence on many interrelated factors (Probst et al., 2020).

## CONCLUSIONS

In this study, we analyzed the distribution of deathbed of ASF-infected wild boars for successive periods, while also accounting for imperfect detection. The epidemiological situation of ASF among wild boar populations seems to have changed into an endemic status and further appears to be getting worse. Although the performance of the existing surveillance system has improved, it is possible to improve the sensitivity based on the results of this study to suppress the further spread of ASF. We believe that the factors identified in this study could be useful for developing and improving risk-based surveillance systems.

## ACKNOWLEDGEMENT

The authors would like to thank Jin A Kim (Daegu Center for Infectious Diseases Control and Prevention) and Kyung-Duk Min (Seoul National University) for their comments on the study.

